# AssayBLAST v2: Major update improving reliability and reporting of the *in silico* analysis of molecular multi-parameter assays

**DOI:** 10.64898/2026.04.27.721032

**Authors:** Tom Eulenfeld, Maximillian Collatz, Sascha D. Braun, Ralf Ehricht

## Abstract

**Introduction:** Accurate *in silico* evaluation of primers and probes is essential for the rational design of molecular multi-parameter assays. We present AssayBLAST v2 to automate and simplify this process for extensive assay designs.

**Results:** A newly integrated strand and proximity check enables precise validation of corresponding oligonucleotides, ensuring correct orientation and spacing for efficient amplification. Based on predicted oligonucleotide interactions, AssayBLAST v2 estimates amplification outcomes, offering a computational benchmark for downstream wet-lab validation and performance correlation. Additionally, the updated software integrates an adaptive BLAST parameter optimization that dynamically scales with database size, thereby improving both analytical sensitivity and computational performance. These improvements are supported by a comparative evaluation against the previous version of AssayBLAST.

**Conclusions:** Collectively, these enhancements streamline the assay development workflow, reduce costs associated with suboptimal primer and probe synthesis, and increase the robustness and reliability of molecular diagnostics and research applications.

## 1 Background

Molecular methods such as PCR, qPCR, isothermal amplification, and DNA microarrays rely on the specific binding of short oligonucleotides to their preferably unique complementary counterpart (Mullis and Faloona, 1987; Southern, 1975; Hassibi et al., 2009). Therefore, the design of molecular multi-parameter assays critically depends on accurate *in silico* evaluation of primers and probes, a task that is often complex and nontrivial. With the release of AssayBLAST, users gained not only a comprehensive assessment of individual oligonucleotides’ performance but also the ability to detect unexpected off-target binding events early in the design process, thereby reducing the risk of unnecessary and costly wet-lab experimentation (Collatz et al., 2025). Importantly, AssayBLAST allows users to build a custom BLAST database (Basic Local Alignment Search Tool, Altschul et al., 1997) composed of intended target sequences and additional genomes or sequence sets representing potential off-target or background sequences. This flexibility enables the user to refine oligonucleotide designs even for assays requiring extreme specificity, such as those employed in bacterial outbreak screening or clinical diagnostics. Because AssayBLAST predictions are highly accurate in practice, for example, achieving a 97.5% concordance with experimental microarray hybridization results in the original publication Collatz et al. (2025), the output provides a solid foundation for optimizing downstream wet-lab protocols to reflect predicted and optimize binding behavior better.

Version 2 of AssayBLAST was rewritten entirely from scratch. Significant enhancements include automatic assignment and classification of nearby oligonucleotides, as well as improved presentation of results. In addition, key BLAST parameters are now dynamically optimized, which improves the tool’s runtime and reliability. These and further modifications are described in detail in the following section. Evaluations against the previous version of AssayBLAST are presented in section 3.

## 2 Implementation

### 2.1 Automatic adjustment of key BLAST parameters

In AssayBLAST, users define match filtering by specifying the number of allowed mismatches (parameter `--mismatch`). BLAST does not allow the direct use of this parameter. Instead, the *E-value* is used. The E-value is the expected number of random matches with the same or better score found in a database of the same size. It is a critical parameter because BLAST filters hits in an early screening step. Using a small E-value results in a much shorter runtime. However, the E-value must not be too small, or it will false negatively miss hits that fulfill the matching criteria. In previous versions of AssayBLAST, the E-value was hard-coded to 1 000 (Collatz et al., 2025). With this setting, however, many, if not all, BLAST hits were missed when AssayBLAST was used to screen a larger database. For instance, for databases larger than 300 MB, hits with two or more mismatches are missed for a query length of 18 nucleotides. Therefore, the E-value was increased to 100 000 for version 1.0. In fact, it became necessary to manually update the E-value depending on the size of the current database. In version 2, the optimal E-value, guaranteeing reliability and best performance, is automatically calculated based on the database size. In a first step, the alignment score *S* for a hit of the shortest query with the maximum number of allowed mismatches is calculated with

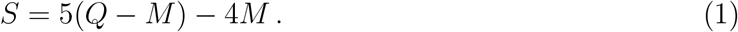

Here, *Q* denotes the length of the shortest query sequence and *M* the number of allowed mismatches (parameter `--mismatch`, default is 2). The reward for a match is 5, the penalty for a mismatch is 4. As gaps in sequence alignments do not occur in biological reality but are merely artifacts of sequence comparison algorithms, we assigned a high gap cost of 1 000 to prevent their inclusion in BLAST alignments. The bit score *S*_2_, the log_2_-scaled required size of a database to get a match by chance, is calculated from the score *S*. The linear relationship between *S*_2_ and *S* depends on the scoring system and the derived parameters of the Gumbel distribution, which are calculated within BLAST (Karlin and Altschul, 1990; Altschul, 1991; Altschul et al., 1997). We approximate this relationship by performing a simple linear regression between the calculated scores *S* and BLAST-reported bit scores *S*_2_, based on hits spanning different primer lengths and numbers of mismatches (see figure 1):

**Figure 1:**
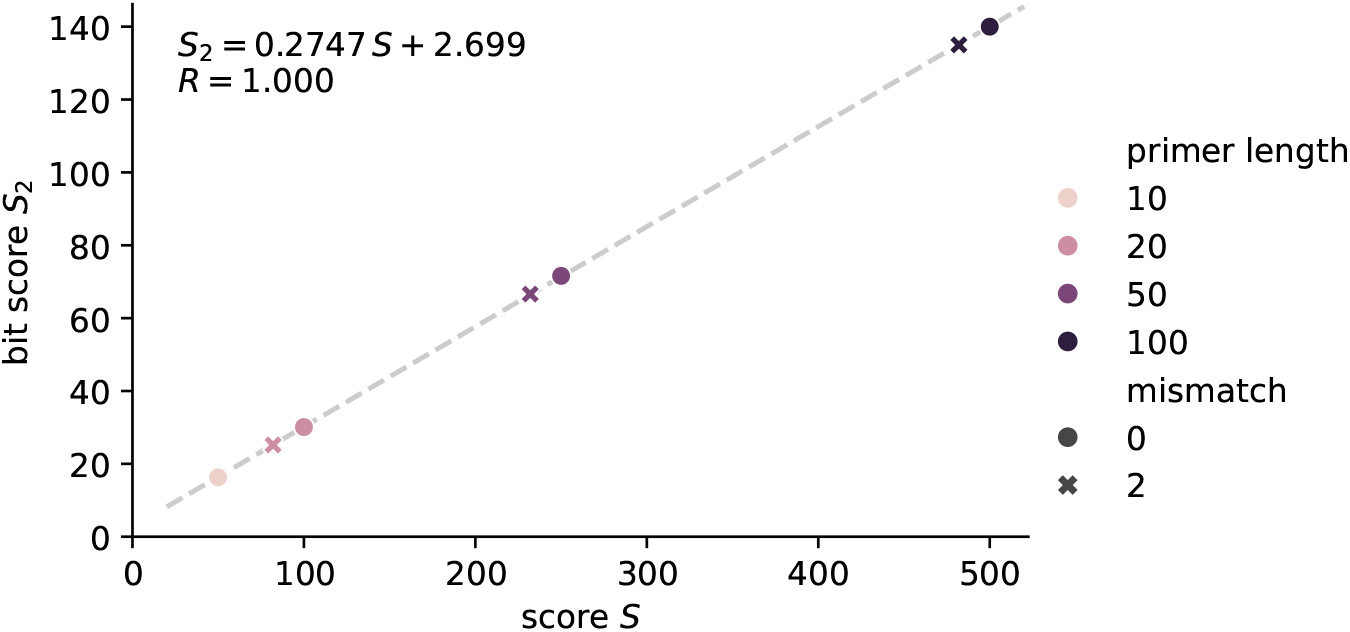
Relationship between BLAST bit score and raw alignment score. The mapping depends on the underlying scoring system, which is fixed in AssayBLAST.

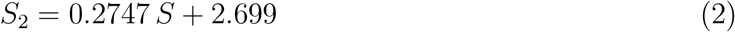

We verified that this relationship is independent of sequence characteristics by evaluating genomes from different organisms. The E-value *E* used in the BLAST call can be calculated from the bit score *S*_2_ and the database size *D* with

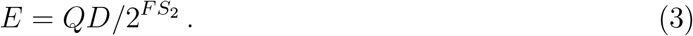

The safety factor, denoted by *F* =0.95, accounts for rounding errors and the internal BLAST recalculation of E-values (González-Pech et al., 2018). Although the calculated E-value guarantees the identification of all matches with the permitted number of nucleotide mismatches or fewer, matches with more mismatches may be included due to queries longer than *Q*. These hits are filtered out later when the BLAST results are analyzed. Finally, the BLAST parameter `-max_target_seqs` is set to 1 000 000 000 to ensure that no hits are missed. It is in the order of the maximum allowed value for this parameter, a signed long integer (2^31^ − 1).

### 2.2 Classification of nearby oligonucleotides and enhanced reporting

Version 1.0 of AssayBLAST reported the BLAST hits in a table with columns for each oligonucleotide-strand combination and rows for each database sequence. Table 1 reports the results of version 1.0 using the dataset of Collatz et al. (2025) reduced to three sequences. In previous versions, users manually paired probe and primer hits based on positional proximity and strand orientation to infer amplification types.

**Table 1:** Obsolete results table for AssayBLAST version 1.0. Each oligonucleotide-strand combination has its own column. Theoretical amplifications are not inferred. The displayed data is an excerpt of the used data (Collatz et al., 2025).

|  | primer_lukF<br>_11b_forward | primer_lukF<br>_11b_revcomp | probe_lukF<br>_10_forward | probe_lukF<br>_10_revcomp | primer_entN<br>_51_forward | primer_entN<br>_51_revcomp | probe_entN<br>_11_forward | probe_entN<br>_11_revcomp |
| --- | --- | --- | --- | --- | --- | --- | --- | --- |
| CP102956 |  | 0 (pos:<br>1942449-1942466) | 0 (pos:<br>1942420-1942445) |  |  |  |  |  |
| CP102957 | 0 (pos:<br>2204112-2204129) |  |  | 0 (pos:<br>2204133-2204158) |  |  |  |  |
| CP102958 | 0 (pos:<br>2344146-2344163) |  |  | 0 (pos:<br>2344167-2344192) |  | 0 (pos:<br>184189-184210) | 0 (pos:<br>184157-184181) |  |

The new version of AssayBLAST performs these tasks automatically and can therefore be used to identify and differentiate designed and unintended oligonucleotide interactions, as exemplified in figure 2. For this purpose, the BLAST hits for each searched sequence are first sorted by their genomic position. Then, the tool iterates through all BLAST hits corresponding to probes and searches for primer hits up to a specified nucleotide distance on the 5’ and 3’ sides (`--distance` option, default is 250 nucleotides). Additionally, the user can switch to a mode that only checks for primer-primer interactions with the `--only-primer` flag. The results are reported in two tables in tab-separated value (TSV) format. The overview table, see table 2, reports the inferred amplification for each probe and sequence, along with a string representing the best probe/primer hits. Table headers correspond to user-defined probe names provided in the input file. Valid amplification values are:

**Table 2:** The AssayBLAST version 2 overview table indicates the inferred amplification and provides a description of the paired oligonucleotides for each probe and database sequence. Table headers correspond to user-defined probe names provided in the input file. Cells use the following abbreviations: F: forward primer; P: probe; R: reverse primer. Further details are provided in the text. The table was generated using the same dataset as in table 1.

| Genome | probe_entN_11 | probe_lukF_10 |
| --- | --- | --- |
| CP102956 | none | lin P0+ R0- |
| CP102957 | none | lin F0+ P0- |
| CP102958 | lin P0+ R0- | lin F0+ P0- |

**Figure 2:**
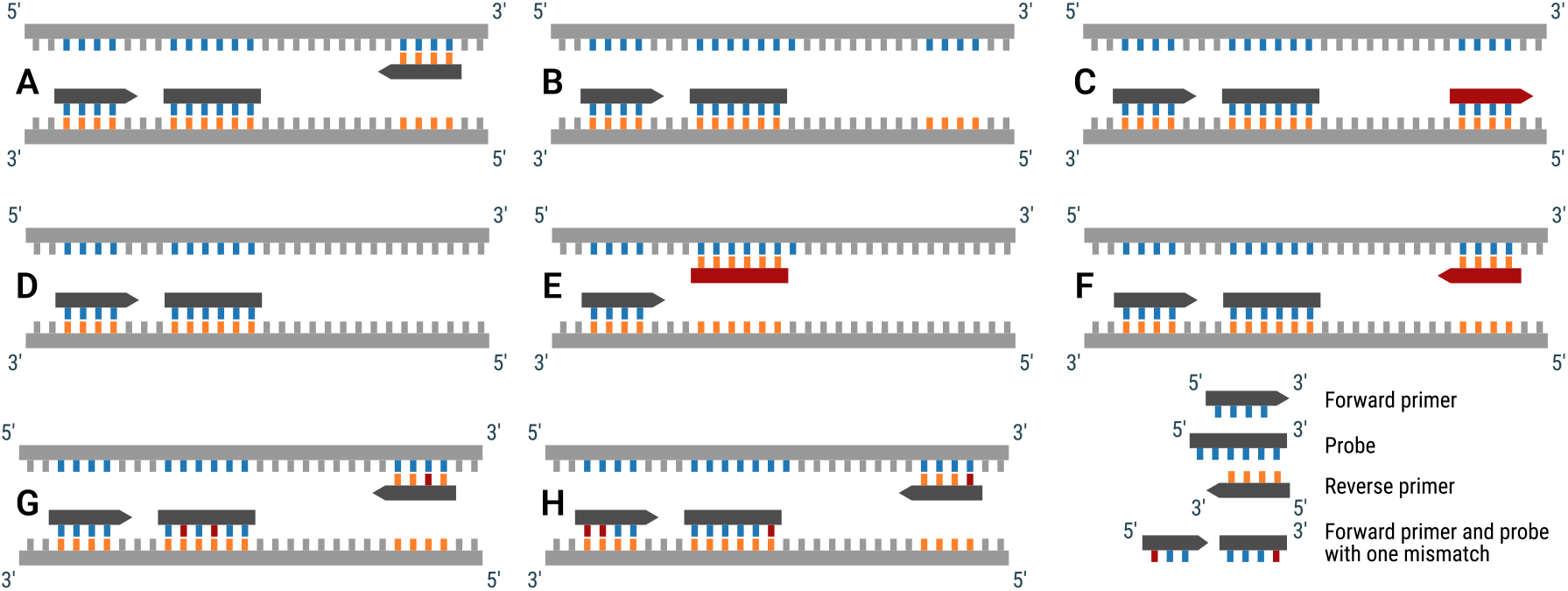
Oligonucleotide interactions identified by AssayBLAST version 2. Top row: A user-designed interaction with exponential amplification is identified in example A. In example B, the reverse primer is missing or binds outside of the specified distance. In example C, the reverse primer binds to the wrong strand. For both examples B and C a linear amplification is reported by AssayBLAST. Middle row: A user-designed interaction with linear amplification is identified in example D. In example E, the probe binds to the wrong strand, leading to no amplification (i.e., constant). In example F, an additional primer—unintended by the user—binds to the forward strand, resulting in unexpected exponential amplification. Bottom row: AssayBLAST also reports the number of mismatches for each oligonucleotide (G), specifically highlighting mismatches at the probe’s 5’ end or the primer’s 3’ end (H), as these negatively affect the transcription.

- `none`: no probe found,
- `const`: probe found, but no primers found,
- `lin`: linear amplification, a working primer was found on one side of the probe, and
- `exp`: exponential amplification, working primers were found on both sides of the probe.

The oligonucleotide combination pattern follows the regular expression:

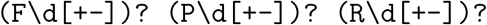

`F, P`, and `R` denote the forward primer (located on the 5’ side of the probe), the probe and the reverse primer (located on the 3’ side of the probe), respectively. The number after that denotes the number of mismatches of the hit, and finally, the strand on which the hit is located is indicated by + or -. Note that a “working” primer necessarily requires the forward primer to be located on the forward strand (+) and the reverse primer to be located on the reverse strand (-).

The detailed table, see table 3, reports a row for each oligonucleotide interaction. Therefore, each cell in table 2 corresponds to a row in table 3. Each row states the genome, the probe ID, the expected amplification, and the paired oligonucleotide IDs and strand directions. The next column gives the strand directions of the oligonucleotides. The strand check column displays the result checking whether the primers are located on the correct strand (see above). The mismatch column displays the number of mismatching nucleotides for each hit. A mismatch directly at the 5’ end of the probe or the 3’ end of the primer is indicated by an asterisk, as this negatively influences their interaction. The distance column displays the distance in nucleotides between the hits. The final column shows the exact locations of the BLAST hits.

**Table 3:** The AssayBLAST version 2 detailed table characterizes all paired oligonucleotide hits. Each cell in table 2 corresponds to a row in table 3. The abbreviations in the column headers are as follows: Amp: Amplification; SD: Strand direction; SC: Strand check; MM: Mismatch; Dist: Distance. The table was generated using the same dataset as in table 1.

| Genome | Probe | Amp | Pairing and Strand direction ((+)/(-)) | SD | SC | MM | Dist | Location |
| --- | --- | --- | --- | --- | --- | --- | --- | --- |
| CP102956 | probe_lukF_10 | lin | probe_lukF_10 (+), primer_lukF_11b (-) | +- | pass | M0,0 | D3 | 1942420:1942445:+,1942449:1942466:- |
| CP102957 | probe_lukF_10 | lin | primer_lukF_11b (+), probe_lukF_10 (-) | +- | pass | M0,0 | D3 | 2204112:2204129:+,2204133:2204158:- |
| CP102958 | probe_entN_11 | lin | probe_entN_11 (+), primer_entN_51 (-) | +- | pass | M0,0 | D7 | 184157:184181:+,184189:184210:- |
| CP102958 | probe_lukF_10 | lin | primer_lukF_11b (+), probe_lukF_10 (-) | +- | pass | M0,0 | D3 | 2344146:2344163:+,2344167:2344192:- |

### 2.3 Other updates

We now publish AssayBLAST on PyPI, which allows for easy installation. Beginning with version 2, the workflow has been modularized into two command-line utilities. The `assay_blast` script calculates the BLAST parameters and runs the BLAST search. The results are analyzed with the `assay_analyze` script. Since the BLAST search is most time-consuming part of the pipeline, this separation enables reuse of stored BLAST results, allowing downstream analyses with modified parameters without re-executing the BLAST search. Additionally, AssayBLAST version 2 uses a single BLAST call for the forward and reverse strands, rather than two separate calls as in version 1.0, since BLAST inherently evaluates both strands within a single query. This consolidation reduces redundant computations and improves runtime efficiency. We also introduced a warning for duplicate primer or probe sequences in the query file, including detection of reverse-complement duplicates. Note that this check does not model partial primer-primer interactions. The BLAST output is parsed using the Sugar library (Eulenfeld, 2025), which supports the detection and parsing of various feature formats. Consequently, the `assay_analyze` script can also be used with MMSeqs2 output (Steinegger and Söding, 2017) or GFF/GTF files. The Sugar library is also used for the oligonucleotide matching.

## 3 Results and Discussion

To confirm the amplification prediction of the updated pipeline, we reproduced the results of the study by Collatz et al. (2025) using the dataset provided in their supplementary material. We use the same threshold of 0.5 to classify microarray experiments as positive or negative. The confusion matrix comparing the automated interpretation of AssayBLAST version 2 results versus the DNA microarray survey is displayed in figure 3. The false positive rate is reduced by more than 50% compared to figure 1 in Collatz et al. (2025), resulting in an increase in accuracy from 97.5% to 98.1%. The improved results may partly be explained by human error during the manual analysis of more than 1 000 positive predictions required prior to AssayBLAST v2.

**Figure 3:**
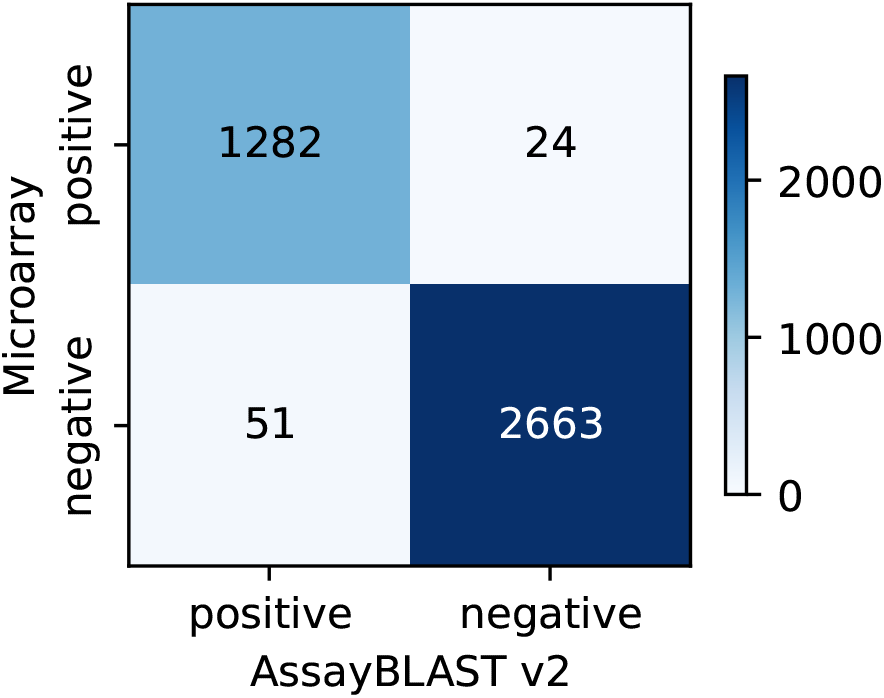
Confusion matrix comparing the automated interpretation of AssayBLAST version 2 results with the DNA microarray results reported in Collatz et al. (2025). The accuracy is improved compared to the previous version of AssayBLAST (Collatz et al., 2025, figure 1).

We also compare the correctness and runtime of AssayBLAST v2 with the previous version. For this purpose, we downloaded complete genomes from diverse organisms: *Influenzavirus A* (H3N2 subtype, `GCA_039834415.1`), *Staphylococcus aureus* (`GCF_000568455.1`), *Candida albicans* (`GCF_000182965.3`), and *Homo sapiens* (T2T, `GCF_009914755.1`). We generated primers of lengths 10 nt, 20 nt, 50 nt, and 100 nt by selecting sequences from the beginning of each genome. We then evaluated different versions of AssayBLAST to identify interactions between each primer and genomic regions. For AssayBLAST version 1, we considered two configurations: one with a fixed E-value of 1 000, as reported in Collatz et al. (2025), and one with a fixed E-value of 100 000, corresponding to the version tagged on GitHub. For the evaluation we use as single core of an AMD EPYC 9754 128-Core Zen 2.25 GHz machine. Correctness of results and run times are reported in table 4. Two issues were identified with AssayBLAST v1. First, AssayBLAST v1 could not correctly identify interactions in cases where the fixed E-value was lower than or comparable to the automatically calculated E-value of version 2. This occurs for combinations of large genomes and short primers, particularly when mismatches are allowed (e.g., table 4, *Candida albicans*, 10 nt). Second, when mismatches are allowed, AssayBLAST v1 inadvertently permits gaps in the genome during BLAST alignment. As a result, erroneous interactions are reported in version 1 (e.g., *Homo sapiens*, 50 nt). Conversely, when the fixed E-value in version 1 is much higher than the automatically calculated E-value, AssayBLAST v2 shows improved runtime compared to the previous version, due to increased BLAST search efficiency. The largest speedup was observed for long genomes and long primers (e.g., *Homo sapiens*, 100 nt).

**Table 4:**
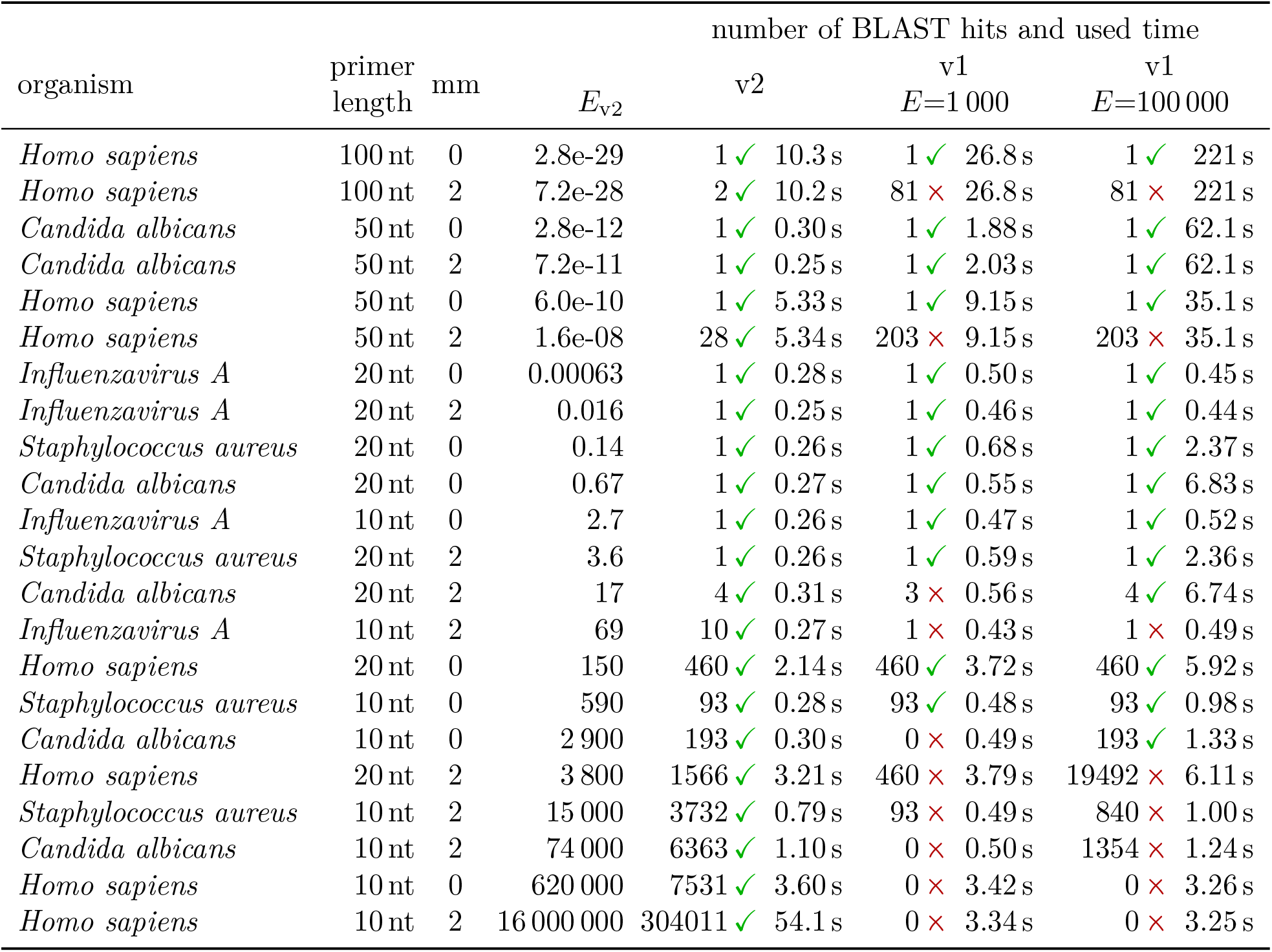
Evaluation of AssayBLAST v2 compared to AssayBLAST v1. The number of BLAST hits is reported for different organisms, primer lengths, and allowed mismatch counts (column mm), along with whether AssayBLAST v1 correctly identified these hits and the corresponding runtime. The automatically determined E-value (column *E*_v2_) improves both analytical sensitivity and computational performance. The table is sorted by this column.

In the original workflow, primer–probe interaction outcomes were derived through manual inspection and interpretation of the AssayBLAST output (table 1). In contrast, AssayBLAST v2 determines these outcomes automatically by applying the described rule set using strand orientation, spatial proximity, and mismatch counts (tables 2 and 3). This transition from manual to automated interpretation improves reproducibility, efficiency, and avoids human error, especially when dealing with large and complex assay designs.

A webservice with a similar perspective is Primer-BLAST (Ye et al., 2012, https://www.ncbi.nlm.nih.gov/tools/primer-blast, last accessed 26/02/2026), which integrates PRIMER3 (Rozen and Skaletsky, 2000) for primer design and enables specificity screening of individual primer pairs against a user-selected reference database. In contrast, AssayBLAST is tailored for the evaluation of multiple primer and probe sets within complex assay configurations and provides a comprehensive assessment of their amplification performance across target sequences. The tool further reports unintended or off-target amplification events. As a command-line application, AssayBLAST ensures full user control and maintains complete data sovereignty. AssayBLAST should be seen as a complementary quality check to web tools like Multiple Primer Analyzer (https://www.thermofisher.com, last accessed 27/10/2025) which check primer properties like melting temperatures and primer-primer interaction, but do not include any screening of databases of interest.

It has been demonstrated that AssayBLAST can accurately predict microarray results. (Collatz et al., 2025). By bridging *in silico* and experimental validation, AssayBLAST 2 supports more reliable, reproducible, and scalable molecular diagnostic and research workflows.

## 4 Conclusions

AssayBLAST 2 formalizes two critical components that should be integral to any modern primer design pipeline. First, it performs *in silico* quality control of designed oligonucleotides, including assessment of target binding efficiency, strand specificity, primer–probe distance verification, detection of duplicate oligonucleotides, and identification of potential non-specific binding events. Second, it provides a prediction of amplification behavior based on the previously calculated oligonucleotide characteristics, serving as a computational reference for comparing wet-lab results and subsequent protocol optimization. These advancements establish AssayBLAST 2 as a robust, reproducible, and computationally efficient framework for oligonucleotide design, minimizing experimental trial-and-error and enhancing confidence in *in silico* predictions.

## Availability and requirements

**Project name** AssayBLAST

**Project homepage** https://github.com/rnajena/AssayBLAST

**Releases** are distributed via PyPI and archived at Zenodo (Eulenfeld and Collatz, 2025)

**Operating systems** Linux, MacOS

**Programming Language** Python

**Other requirements** BLAST, Python 3.11+, rnajena-sugar

**License** MIT

**Restrictions to use by non-academics** MIT license

## Declarations

### Ethics approval and consent to participate

Not applicable.

### Consent for publication

Not applicable.

### Availability of data and materials

AssayBLAST is available on GitHub at https://github.com/rnajena/AssayBLAST and archived on Zenodo (Eulenfeld and Collatz, 2025). Code to reproduce the results is available at https://www.github.com/rnajena/assayblast_paper (Eulenfeld, 2026). The used genomes can be downloaded from NCBI (https://www.ncbi.nlm.nih.gov). The corresponding accession numbers are provided in Eulenfeld (2026).

### Competing interests

The authors declare that they have no competing interests.

### Funding

This research was funded by the Deutsche Forschungsgemeinschaft (DFG, German Research Foundation) under Germany’s Excellence Strategy – EXC 2051, Project-ID 390713860.

### Author contributions

R.E. conceptualized the tool. T.E. and M.C. developed AssayBLAST version 2. S.B. tested the software. T.E. and M.C. wrote the manuscript. All authors revised the manuscript.

## Acknowledgments

The authors thank the anonymous reviewers for their insightful and constructive comments.

## Notes

### Competing Interest Statement

The authors have declared no competing interest.

